# FLEX-DBS: a flexible wireless platform for bilateral constant-current deep brain stimulation in freely moving rodents

**DOI:** 10.64898/2026.10.09.757775

**Authors:** Robert D. Graham, Meaghan C. Creed, Matt Gaidica

## Abstract

**Background:** Preclinical deep brain stimulation (DBS) in rodents is central to understanding how DBS works, but most rodent DBS is delivered through a tether, which constrains behavior, home-cage experiments and uninterrupted stimulation. Most long-runtime wireless mouse stimulators achieve their runtimes by setting stimulation parameters before or around implantation and operating from supplies of about 3 V.

**New method:** FLEX-DBS (Flexible Deep Brain Stimulation) is a head-mounted, battery-powered stimulator that delivers bilateral constant-current stimulation from improved Howland current pumps on ±5 V rails. Amplitude, pulse width and frequency are programmable in continuous and burst modes from an iOS application over Bluetooth Low Energy, and stimulation continues autonomously after the phone disconnects. The device measures 19.5 × 22.5 × 8.72 mm, weighs 3.71 g in operation and runs on two size-13 zinc-air cells, which are exchanged by briefly detaching the device, pausing stimulation only for the exchange.

**Results:** Into a 1 kΩ bench load, delivered current increased approximately linearly with the amplitude setting and was repeatable from pulse to pulse, with an asymmetric biphasic waveform whose recovery phase ran at about one fifth of the stimulating amplitude. Battery current was 1.47–1.89 mA in standby and 3.15–3.42 mA during stimulation at amplitude settings of 60–400 µA, giving a projected three days of continuous stimulation per pair of cells. Firmware v0.5 left a net charge of 26–35 % of the stimulating-phase charge per pulse into 1 kΩ. Into 47 kΩ, within the in vivo load range, stimulating current was compliance-limited at 93 µA. Pulse-derived estimates of the effective load presented by five implanted platinum electrodes in three mice had weekly means of 43–54 kΩ over eight weeks (32–80 kΩ across individual sessions), placing the maximum regulated current near 90 µA (81–101 µA at the weekly means). Analysis of the current-pump network bounds load-dependent current error at 3–8 % for these loads. In vivo, we delivered cingulate FLEX-DBS at high amplitude settings intended to evoke a robust stimulation-associated behavioral phenotype in six mice. Time spent immobile in an open field rose from 46.5 % at baseline to 75.7 % during stimulation (a 62.7 % increase in the group mean) and remained elevated after stimulation ended; without a sham control, this increase cannot be separated from within-session habituation.

**Comparison with existing methods:** Among the published wireless rodent stimulators compared here, several offer some of these capabilities, but none combines bilateral constant-current output, real-time wireless control that continues as autonomous stimulation after disconnect, and both continuous and burst modes; FLEX-DBS does so at the cost of runtime measured in days rather than weeks.

**Conclusions:** FLEX-DBS is designed for untethered DBS experiments in which parameters must be set, changed and verified at the cage. Its measured power consumption permits a projected three days of continuous operation per cell pair; firmware v0.5 requires improved charge balancing before prolonged in vivo stimulation. The electrode load, rather than the programmable range, sets the usable stimulation amplitude in vivo. In vivo, stimulation at high amplitude settings was associated with increased immobility, consistent with effective stimulation through implanted electrodes; electrode impedance and delivered current were not measured in these mice.

## 1. Introduction

Deep brain stimulation (DBS) is a surgical therapy in which electrical current is delivered to discrete brain nuclei through implanted electrodes (Lozano et al., 2019). Although it is established primarily as a treatment for movement disorders, DBS is increasingly applied to psychiatric indications, chronic pain, neurodegenerative disorders and developmental disorders (Lozano et al., 2019; Krauss et al., 2021; Knotkova et al., 2021). Despite this growing adoption, the neural mechanisms underlying the therapeutic effects of DBS remain poorly understood, which limits both the rational optimization of stimulation parameters and patient selection (Murphy et al., 2021). Preclinical research in rodents therefore has an important role in determining the behavioral, circuit and synaptic mechanisms through which DBS acts (Creed et al., 2015; Creed, 2018).

Preclinical DBS is commonly delivered with the rodent physically tethered to an external pulse generator (Ewing et al., 2013), which constrains experimental design in several ways. A tether restricts the animal’s movement and the environments in which it can be tested (Jiang et al., 2017); its measurable effects on stress markers and sleep can be small (Aulehner et al., 2022), but the restriction itself remains. These consequences complicate or prevent behavioral assays that depend on unconstrained motor behavior, such as spontaneous pain behaviors, movement-disorder phenotypes and social interaction, and so limit the translational relevance of behavioral observations. Tethering is also difficult to implement in the home cage, so stimulation is usually delivered in dedicated apparatus such as open-field chambers, which can itself alter naturalistic behavior. Together with the cost of conventional stimulators, these constraints limit the throughput and statistical power of preclinical DBS studies (Ewing et al., 2013).

Tethering also limits the timescales over which DBS can be studied. Clinically, DBS is delivered continuously for months to years, and its efficacy can wane over time for reasons that remain unclear (Fasano and Helmich, 2019). Some effects of DBS also develop over longer periods than the acute motor effects that dominate preclinical work: pallidal DBS for dystonia produces a delayed, progressive improvement over months (Kupsch et al., 2006), and in rats the behavioral effect of repeated subthalamic stimulation took several sessions to emerge (Aleksandrova et al., 2013). A wireless, battery-powered system for delivering sustained DBS in rodents would support mechanistic studies of both long-term loss of efficacy and indications in which therapeutic effects emerge slowly.

The few wireless rodent stimulators that exist were largely designed for chronic implantation. Most set the stimulation waveform and parameters before implantation or allow changes only through a near-field or cable programmer, which makes it difficult to change parameters during an experiment or to adjust them as the impedance of the electrode–tissue interface changes over the life of the implant. This is a critical limitation because stimulation parameters that produce a behavioral effect in one paradigm can be inert or even opposite in another, and the most informative preclinical studies have depended on varying frequency, amplitude and target within an experiment (Murphy et al., 2021). Low-frequency stimulation of the nucleus accumbens combined with pharmacology, for example, reversed cocaine-evoked synaptic plasticity where conventional high-frequency stimulation did not (Creed et al., 2015), and the effects of subthalamic stimulation on reward-related behavior, relevant to its motivational side effects, depend on stimulation parameters and task design (Vachez and Creed, 2020). Most published long-runtime mouse-scale systems also run from supplies of about 3 V (Paulat et al., 2026; Grotemeyer et al., 2024; Fleischer et al., 2020; Alpaugh et al., 2019; Kouzani et al., 2013), and several are single-channel (Grotemeyer et al., 2024; Fleischer et al., 2020; Alpaugh et al., 2019; Kouzani et al., 2013).

An ideal preclinical DBS device would let the experimenter set and change parameters in a freely moving mouse in its home cage, deliver true constant current into electrodes whose impedance is uncertain and changes after implantation, stimulate bilaterally, and support multiday experiments in which a battery exchange interrupts stimulation only briefly. Here we describe FLEX-DBS (Flexible Deep Brain Stimulation), a flexible wireless platform for bilateral constant-current deep brain stimulation in freely moving rodents. FLEX-DBS is a battery-powered, head-mounted device with ±5 V compliance rails; programmable amplitude, pulse width and frequency; continuous and burst stimulation modes; and a Bluetooth Low Energy (BLE) interface to an iOS application. We present the design as one point in a general DBS architecture, characterize the electrode load it must drive in vivo, describe its electrical validation, and compare it with published wireless systems. The design gives compliance, current precision and real-time wireless control priority over the quiescent current needed for maintenance-free implantation. By combining untethered operation over hours to days with parameter flexibility and regulated current delivery, FLEX-DBS is designed to enable home-cage, social and extended-duration DBS experiments that are impractical with tethered stimulators or fixed-parameter implants, and to support systematic preclinical testing of stimulation paradigms for new clinical indications.

## 2. System architecture and implementation

### 2.1 General DBS architecture

Figure 1 presents the stimulator as a set of functional blocks, and Table 1 lists the FLEX-DBS implementation of each: energy source, power conditioning, timing and waveform generation, current regulation, polarity and output routing, charge recovery, supervisory logic, and wireless control. Published rodent stimulators differ mainly in how they implement these blocks, and those choices define the principal trade-offs: battery-direct (typically *∼*3 V) versus boosted rails, constant-current versus constant-voltage output, single versus split supplies, active versus passive charge recovery, real-time wireless control versus autonomous operation, and experimental flexibility versus power optimization for long-term implantation.

**Figure 1.**
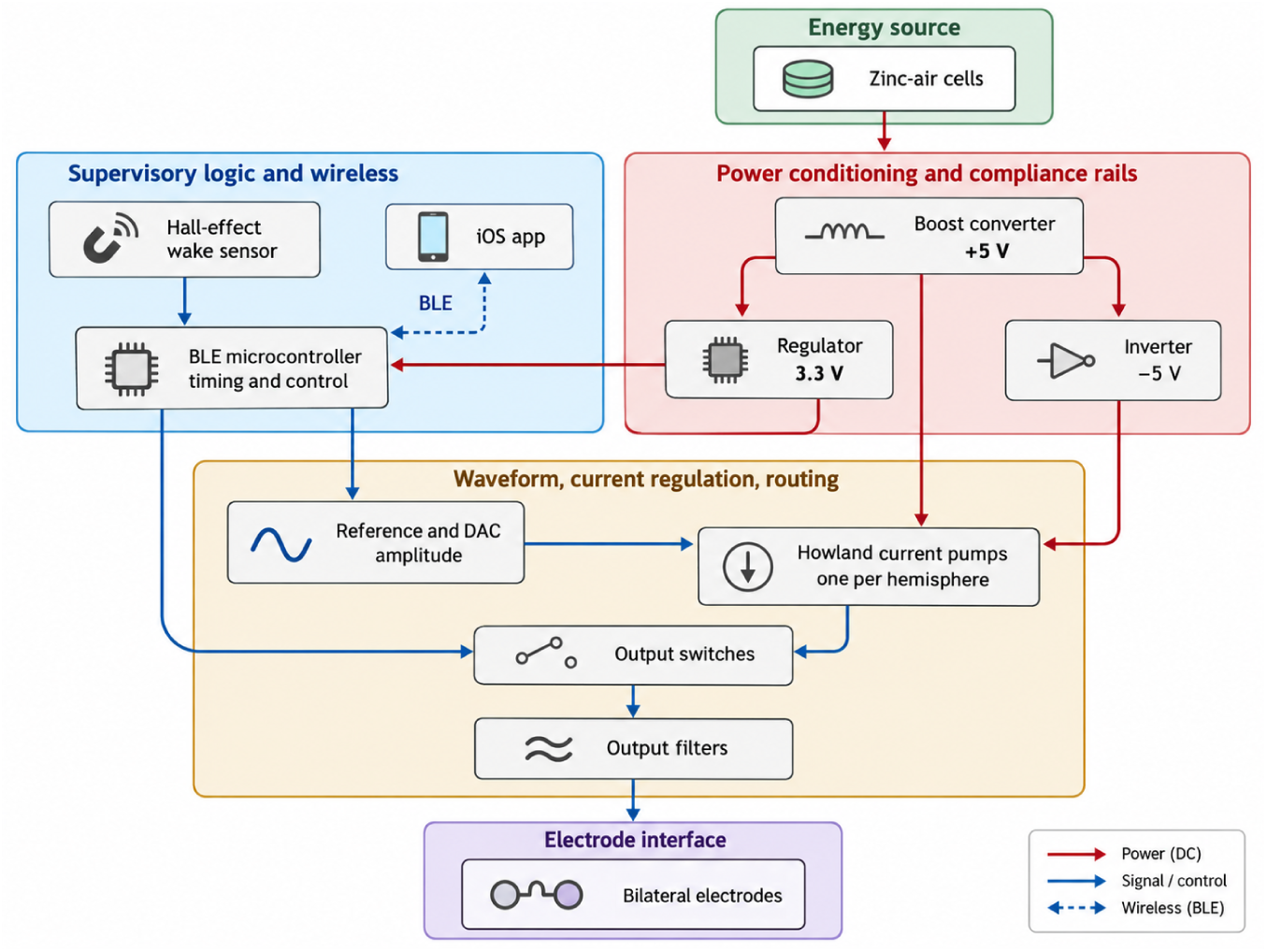
FLEX-DBS architecture. Functional blocks used to describe rodent stimulators, as implemented in FLEX-DBS. Zinc-air cells feed a boost converter that generates the +5 V rail; an inverter derives −5 V, and a regulator derives 3.3 V for the BLE microcontroller. The microcontroller sets pulse timing, commands amplitude through a reference and DAC, and controls the output switches. The DAC drives two Howland current pumps on ±5 V, one per hemisphere, whose outputs pass through the switches and output filters to bilateral electrodes, one per hemisphere, each a twisted pair of a stimulating wire and a ground wire. A Hall-effect sensor wakes the device when a magnet is passed over it, and parameters are set from an iOS application over BLE. Components are listed in Table 1.

**Table 1.** FLEX-DBS implementation of each architectural block.

| Block | FLEX-DBS implementation |
| --- | --- |
| Energy source | Two size-13 (PR48) zinc-air cells in series: 2.8 V nominal, $\sim 280$ mAh |
| Power conditioning | MAX17220 boost converter to +5 V; MAX1853 charge-pump inverter to $-5$ V; TPS709 LDO from +5 V to 3.3 V for logic |
| Timing and waveform | BGM220S microcontroller; pulse edges from a 32.768 kHz low-frequency timer; amplitude from a 16-bit DAC (AD5683) referenced to a 1.25 V precision reference (LT6656) |
| Current regulation | Two improved Howland current pumps (LT6020-1 dual precision op amp) on $\pm 5$ V, one per hemisphere, sharing one DAC command |
| Polarity and routing | ADG1236 dual SPDT analog switch: each pump output either to its electrode or to ground; polarity reversal by stepping the DAC across the 1.25 V reference |
| Charge recovery | Active, asymmetric: reverse-polarity phase at one-fifth amplitude for five times the pulse width |
| Supervisory logic | Same BGM220S; parameters held in non-volatile memory; battery voltage measured by the on-chip ADC |
| Wireless | BLE 5 with a custom text protocol; iOS application (MouseCap) for configuration; Hall-effect sensor (DRV5032) for magnet wake |

### 2.2 FLEX-DBS implementation

FLEX-DBS implements the general architecture with the choices listed in Table 1. The circuit schematic (Supplementary material S1) is available on request, and the firmware, application and communication protocol are released as open source (S2–S4); the main text describes function.

#### Power tree

The two zinc-air cells feed the boost converter, which generates the +5 V rail. The negative rail is derived from +5 V by a charge-pump inverter that is enabled whenever +5 V is present. Logic runs from a 3.3 V linear regulator fed by +5 V. The output stage therefore has a 10 V supply span, and every subsystem, including the microcontroller and radio, sits downstream of the boost converter.

#### Current source

The 1.25 V reference serves as the analog mid-rail and as the external reference of the 16-bit DAC (external-reference variant, gain 2), so the DAC output spans 0–2.5 V centered on 1.25 V. Each Howland pump uses matched 10 kΩ input and feedback resistors on its inverting side and a 12 kΩ reference resistor balanced against a 10 kΩ resistor plus a 2 kΩ sense resistor on its non-inverting side, which satisfies the balance condition of the improved Howland topology (Texas Instruments, 2008). All network resistors are 0.1 % tolerance. At the reference voltage the commanded current is zero; stepping the DAC below the reference produces one polarity and stepping above it produces the other. The ideal transconductance is set by the 2 kΩ sense resistor, 0.5 mA/V, or 6.25 µA per percent of full scale (625 µA at 100 %). The application labels amplitude as a percentage and as the nominal current into a 1 kΩ load, *∼*6 µA per percent (300 µA at 50 % and 600 µA at 100 %).

#### Channels

The device has two output channels, one per hemisphere, each with its own Howland pump, switch and two-stage output filter (two 330 Ω series resistors, each followed by 470 pF to ground; 660 Ω in total per channel). The two channels share the DAC command, so they deliver the same amplitude and, in the current firmware, the same timing: both switches close and open together and both pumps see the same DAC steps. Independent timing per channel is possible with the existing hardware because each switch has its own control line; independent amplitude is not, because the DAC is shared. The four-pin electrode connector carries the two channel outputs and the ground return. One electrode is implanted per hemisphere, and each electrode is a twisted pair of one stimulating wire, connected to the channel output, and one ground wire, connected to the device ground; no separate return electrode is used.

#### Pulse generation

Each pulse is sequenced by the microcontroller as a fixed chain: the op amp is brought out of shutdown and allowed 150 µs to settle with the DAC at the zero-current reference; the switches then connect both pumps to their electrodes and are allowed 30 µs; the DAC steps to the commanded amplitude *I* for the programmed pulse width *P*; the DAC steps to the opposite polarity at *I*/5 for 5·*P*; the DAC returns to the reference; the switches disconnect the electrodes; and the op amp is shut down. Switches change state only at zero commanded current, so the load is never connected to a saturated open-circuit current source and never disconnected mid-pulse, and pulse edges are produced by DAC steps into an already-connected load. The recovery phase is designed to carry the same charge as the stimulating phase; with firmware v0.5 the measured phases leave a net residual in the stimulating direction (Section 4.1). Because the op amp is enabled only around each pulse, its on-time is roughly 210 µs + 6·*P*, about 10 % of the period for a 90 µs setting at 130 Hz and up to about 50 % at 600 µs.

Pulse and inter-pulse timing derive from the 32.768 kHz low-frequency timer, so every duration is quantized to one 30.5 µs tick. To compensate for interrupt latency, tick quantization and the DAC write, the firmware subtracts a fixed overhead estimate from each phase (117 µs from the stimulating phase and 116 µs from the recovery phase). The compensation is not exact: at the 180 µs setting the stimulating phase measured *∼*240 µs and the recovery phase *∼*860 µs rather than 900 µs. The stimulation period is set by a periodic timer whose period is computed in whole milliseconds, rounded down, so the delivered frequency takes one of seven values between *∼*83 and *∼*167 Hz (Table 2). The 130 Hz setting used throughout this work delivers 142.5 Hz; the 100 and 125 Hz settings correspond to whole-millisecond periods and are not rounded. In burst mode a second periodic timer starts a train of pulses at the intra-burst frequency every burst period and stops scheduling new pulses at the end of the burst duration. The first pulse of each burst occurs one intra-burst period after burst onset, and a pulse already in progress when the burst ends completes its full stimulating and recovery chain rather than being truncated.

**Table 2.** Programmable stimulation parameters: application settings and delivered values (firmware v0.5). Measured values are from Section 4.1.

| Parameter | App setting | Delivered | Notes |
| --- | --- | --- | --- |
| Amplitude | 0–100 % in 1 % steps; labeled 0–600 $\mu\text{A}$ into 1 kΩ | 16-bit DAC; ideal transconductance 6.25 $\mu\text{A}$ per percent (0–625 $\mu\text{A}$); into 1 kΩ, $\sim 4.7\ \mu\text{A}$ per percent + 47 $\mu\text{A}$ measured; limited by compliance at high load | Shared by both channels |
| Pulse width $P$ | 90–600 $\mu\text{s}$ in 30 $\mu\text{s}$ steps | Quantized to 30.5 $\mu\text{s}$; 238–254 $\mu\text{s}$ measured at the 180 $\mu\text{s}$ setting | Recovery phase set to $5 \cdot P$ at $I/5$ |
| Frequency (continuous mode) | 80–160 Hz in 5 Hz steps | Period in whole milliseconds, rounded down: $\sim 83, 91, 100, 111, 125, 143$ or 167 Hz | 130 Hz setting delivers 142.5 Hz (measured) |
| Burst period | 1 ms – 60 min | As set | Burst mode only |
| Intra-burst frequency | 80–160 Hz in 5 Hz steps | Same seven values as continuous mode | Burst mode only |
| Burst duration | 1 ms – 120 s | Capped at the burst period; about duration ÷ intra-burst period pulses per burst | Burst mode only; in-flight pulse completes |
| Device ID | 00–99 | — | Displayed in the app for multi-animal cohorts |

#### Parameters

Table 2 lists the parameters exposed by the application and what the hardware and firmware v0.5 deliver for each. Parameters are stored in the microcontroller’s non-volatile memory and survive power cycling, so a device configured in the laboratory delivers the same protocol after its cells are replaced in the behavior room or home cage.

#### Wireless control and user interface

The device advertises as a BLE peripheral at a 2 s interval and exposes one service with a write characteristic (app to device) and a notify characteristic (device to app). Commands and responses are short ASCII strings of comma-separated key–value pairs; for example, `_M0,A50,F130,P90,G0,N0` sets continuous mode, 50 % amplitude, a 130 Hz frequency setting, a 90 µs pulse-width setting, activation off and device number 0. The device replies with its full configuration, battery voltage and firmware version. The negotiated ATT MTU (up to 247 bytes) allows a complete configuration in a single packet. The iOS application (Figure 3) presents amplitude, pulse width and frequency as sliders with the derived current shown alongside, a segmented control for continuous versus burst mode, and a toggle that starts the stimulation protocol. A red outline on the Sync button indicates that on-screen values differ from those confirmed by the device.

The intended workflow is configure-then-run: the experimenter connects, sets parameters, arms activation and disconnects; the device begins stimulating and stops advertising. Stimulation continues autonomously with no radio link. Passing a magnet over the device’s Hall-effect sensor re-enables advertising while stimulation continues; the phone can then reconnect to inspect battery voltage, change parameters without interrupting stimulation, or stop it, all without handling the animal. A status LED indicates when the device is advertising or connected and turns off during stimulation.

#### Battery telemetry

The microcontroller samples the battery voltage on request and reports it in millivolts; the application maps 1.4–2.8 V to a 0–100 % indicator.

#### Physical design

The circuit is built on a 1 mm thick printed circuit board measuring 19.5 mm × 22.5 mm, excluding the programming tab, and the assembled device is 8.72 mm thick (Figure 2). Firmware is loaded through a Tag-Connect TC2050 footprint on a break-away programming tab (Figure 2A, P), which can be cut off once firmware is loaded. Silicon Labs over-the-air device firmware update (DFU) is not currently enabled but could be added to allow firmware updates without the tab. A custom snap-on shroud covers the device (Figure 2B). The shroud was designed in Fusion 360 (Autodesk; design available at https://a360.co/3TqHAxf) and printed in PLA on a Bambu Lab X1 Carbon printer. The device with its electrode connector weighs 1.61 g with the programming tab removed (2.17 g with the tab), the shroud 0.48 g and each zinc-air cell 0.81 g, for 3.71 g in operation with two cells and the shroud (3.23 g without the shroud).

**Figure 2.**
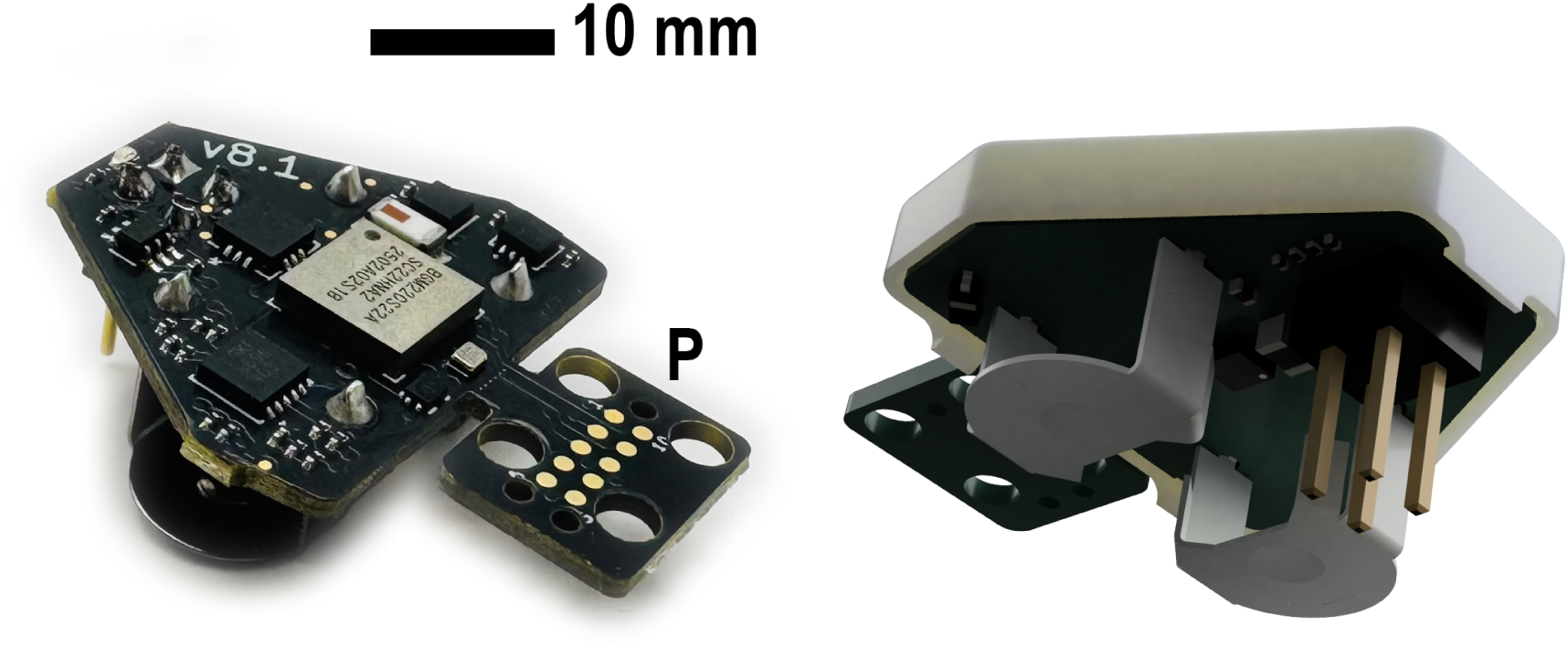
FLEX-DBS hardware. *(A)* Assembled device viewed from above; scale bar, 10 mm. P, break-away programming tab carrying the Tag-Connect TC2050 footprint, which can be cut off after firmware is loaded. *(B)* Rendering of the device from below with the snap-on shroud, showing the two zinc-air cell holders and the electrode connector. The board measures 19.5 mm × 22.5 mm excluding the programming tab, and the assembled device is 8.72 mm thick.

**Figure 3.**
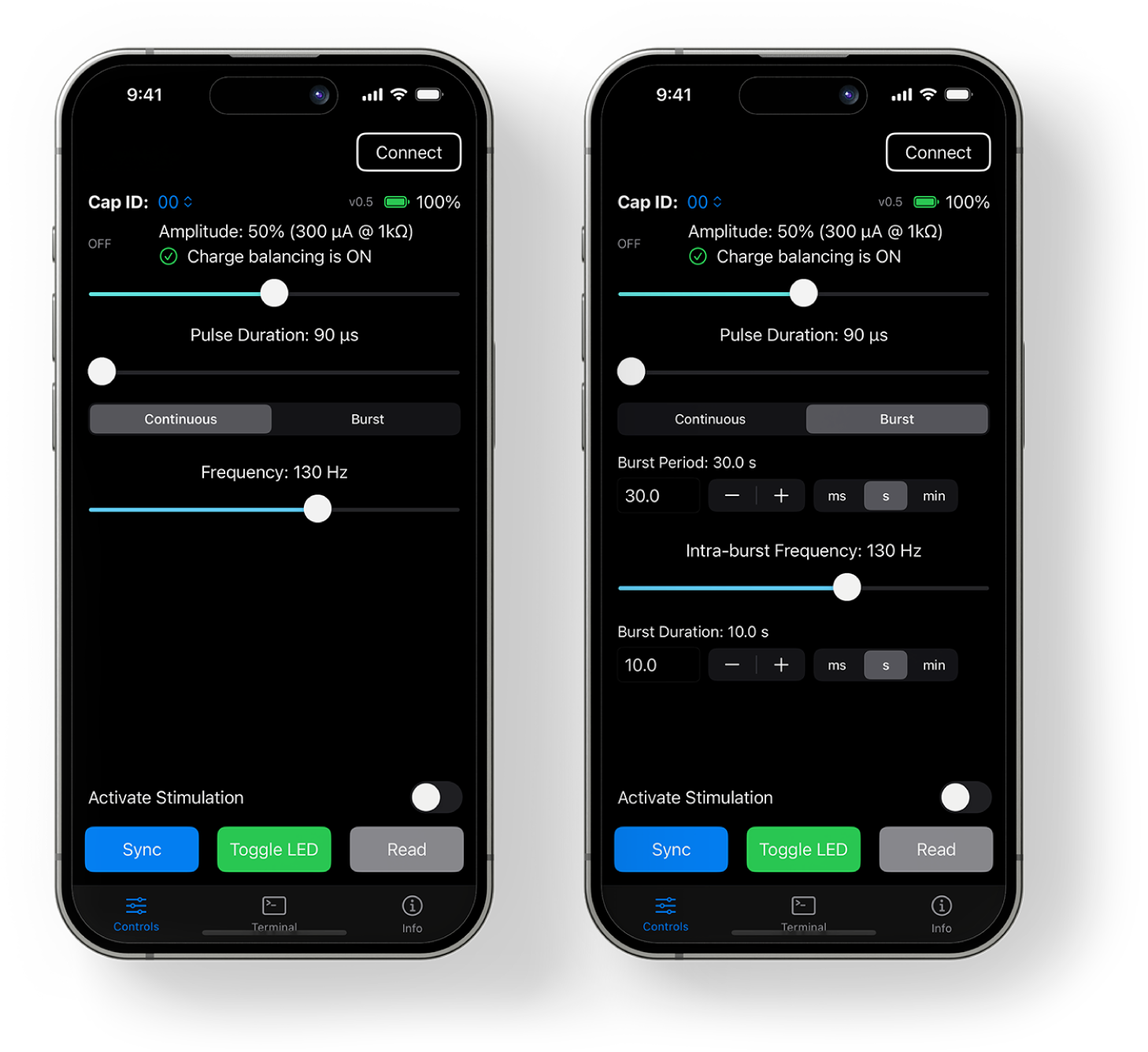
MouseCap iOS application. *(Left)* Continuous mode: amplitude (percent of full scale and nominal current into 1 kΩ), pulse duration and frequency sliders; device ID, firmware version and battery indicator; activation toggle, Sync, LED and Read controls. *(Right)* Burst mode: burst period, intra-burst frequency and burst duration with unit selection. Representative settings are shown (50 %, 90 µs, 130 Hz; 30 s burst period, 10 s burst duration). Displayed values are settings; delivered values are given in Table 2.

### 2.3 Design considerations

The architecture follows from a set of coupled experimental requirements rather than from a single optimum.

#### Stimulation envelope

Parameter ranges were chosen to reproduce protocols with established behavioral effects in our prior rodent DBS work rather than from theoretical models of neural activation. Stimulation at 100 µA, 90 µs and 130 Hz (Creed et al., 2012, 2013) is the reference protocol around which the application’s defaults are written. The frequency range covers the 130 Hz clinical standard and the 80–160 Hz range selected for this device; it does not extend to the 30 and 60 Hz conditions used in earlier comparative work (Creed et al., 2011). Amplitude is programmable in *∼*6 µA steps to 600 µA nominal. Our prior protocols used 100 µA (Creed et al., 2012, 2013) and, in one behavioral study, 12.5 µA (Aleksandrova et al., 2013); the range above 100 µA provides headroom for titration and for lower-impedance electrodes. Firmware v0.5 does not reproduce the programmed timing exactly, and the amplitude setting is a reproducible command rather than a calibrated delivered current. In practice we titrate amplitude in each animal against its behavioral effect, because electrode impedance and implant location vary between animals; on-board current or impedance sensing would make that titration more direct.

#### Compliance and electrode impedance

A constant-current source needs an output voltage of *I · |Z|* plus headroom for the sense resistor, the output filter and the amplifier’s swing limit. With ±5 V rails, a 2 kΩ sense resistor and 660 Ω of series filter resistance per channel, the voltage available at the electrode in the stimulating phase is approximately 4.3 V at 100 µA and 3.5 V at 400 µA, from the 4.6 V swing measured into 47 kΩ (Section 4.1). The implanted electrodes used here present 32–80 kΩ in vivo (Section 3), so compliance rather than the programmable range sets the largest current that can be delivered. A battery-direct design with *∼*3 V of compliance would support substantially less current into the same electrodes.

#### Load dependence below compliance

An improved Howland pump has infinite output impedance only when its resistor ratios are exactly balanced; any mismatch leaves a finite output impedance *R*_out_ in parallel with the load, so the current delivered into a load *R_L_* falls short of the commanded current by the fraction *R_L_*/(*R_L_* + *R*_out_) (Tucker et al., 2013). We computed *R*_out_ for the FLEX-DBS network (10 kΩ input and feedback resistors, 12 kΩ reference resistor, 10 kΩ feedback resistor and 2 kΩ sense resistor; Section 2.2) at every combination of ±0.1 % resistor tolerance, assuming an ideal op amp. The worst case is about 0.9 MΩ. Into the 32–80 kΩ electrode loads measured in Section 3, plus 660 Ω of output filter resistance, this corresponds to a current shortfall of at most 3–8 %; with nominal resistor values the shortfall is zero. Finite op-amp gain would lower *R*_out_ further, so these values are a bound for the resistor network alone, not a measurement.

#### Waveform and charge balance

Reversing the current through the electrode after each stimulating phase prevents net charge accumulation at the electrode–tissue interface and limits the generation of harmful electrochemical reaction products (Merrill et al., 2005). FLEX-DBS does this actively but asymmetrically: the recovery phase is one-fifth the amplitude for five times the duration. Compared with a symmetric biphasic pulse, the low-amplitude recovery phase is less likely to be excitatory itself and avoids a sharp reversal transient; compared with passive electrode shorting, it recovers charge in a defined time regardless of the electrode’s RC time constant. The ratio is fixed in firmware.

#### Bilateral operation

Bilateral stimulation is standard in our behavioral protocols (Aleksandrova et al., 2013) and is required to reproduce them. The shared-DAC design halves the analog cost of a second channel and guarantees identical amplitude in both hemispheres. It cannot deliver different amplitudes to the two sides; that would require a second DAC or time-multiplexing of the existing one.

#### Wireless interaction

In the configure-then-run model the radio is not part of the stimulation path, so a dropped connection does not interrupt stimulation; the advertising interval, connection behavior and text protocol were chosen for reliability and ease of debugging rather than power.

#### Operating duration versus long-term implant power budget

The boost converter chain, the negative rail, the precision reference and an awake microcontroller were expected to dominate the platform current, and the design accepts that overhead to provide compliance, precision and reconfigurability.

## 3. Electrode impedance and stimulation robustness

A constant-current stimulator regulates current only while the voltage the electrode requires stays within its compliance, so the electrode load determines how much of the programmable range is usable.

### 3.1 Animals and electrode implantation

Adult (≥6 weeks) male and female C57BL/6J mice were sourced or bred from Jackson Laboratory lines and used for all experiments. Mice were group housed (2–5 per cage, single sex) under controlled humidity, temperature and photoperiod (12:12; light onset/offset at 07:00 and 19:00) conditions with ad libitum access to food and water. Anesthesia was induced and maintained with isoflurane (5 % and 1.5 %, respectively). Following craniotomy, DBS electrodes were implanted into the cingulate cortex (coordinates relative to bregma: anteroposterior (AP), −0.6 mm; mediolateral (ML), ±0.3 mm; dorsoventral (DV), −1.25 mm). Animals recovered for 7–10 days before behavioral testing. All procedures were approved by the Institutional Animal Care and Use Committee at Washington University in St. Louis (protocol 24-0211) and conducted in accordance with the Guide for the Care and Use of Laboratory Animals (NIH).

Electrodes were Teflon-insulated platinum twisted-wire pairs (Protech International Inc.; length 10 mm; wire diameter 0.25 mm; diameter with insulation 0.33 mm). The mating end of each wire pair was inserted into a custom 3D-printed electrode holder, designed in Fusion 360 (Autodesk; design available at https://a360.co/4rEBmq2) and printed in clear biocompatible resin, and superglue was applied to the base of each wire to hold the pairs in place. After the glue dried, each wire was cut to the length needed to reach the implantation target, and the ends of each electrode were wet-sanded (3000 grit) to maximize contact surface area. One electrode was implanted per hemisphere. Each electrode was a twisted pair in which one wire was the stimulating contact and the other was connected to ground; no separate return electrode was implanted.

### 3.2 In vivo impedance measurement

Platinum electrodes were implanted bilaterally, one per hemisphere, in four mice across two surgical cohorts (eight electrodes); these mice were separate from those used in Section 6. Impedance was measured weekly from 1 to 8 weeks after implantation. At each session a StimJim stimulator (Cermak et al., 2019) delivered 10 µA, 450 µs pulses at 130 Hz for 10 s through the stimulating wire of each electrode, with the other wire of the pair as ground, and the pulse voltage and delivered current reported by the StimJim were recorded five times. Impedance was computed as voltage divided by delivered current and averaged over the five readings; this quantity is an effective, pulse-derived estimate of the load, referred to as impedance in this section. This two-electrode configuration is the one the stimulator itself drives. Two- and three-electrode impedance spectroscopy measurements are highly correlated (Koo and Weiland, 2018), which supports the two-electrode configuration but does not validate the time-domain pulse estimate used here specifically. This pulse-based estimate matches the quantity that matters for a constant-current stimulator, the voltage required per unit of delivered current, rather than a small-signal impedance at a single frequency.

Three electrodes were excluded. One read near 0 V at every session, consistent with a short between the stimulating and ground wires of the pair. Both electrodes of one mouse gave unstable readings, varying by up to 4.7-fold between sessions and drifting within a session; whether this reflected the electrodes, the implant strategy or the recording setup could not be determined. The analyzed data set therefore comprises five electrodes from three mice, measured at eight weekly sessions (40 electrode-sessions).

Two properties of the measurement bound its interpretation. First, in a bench check against resistive loads, the voltage reported by the StimJim fell 6–73 % below the oscilloscope-measured voltage for loads of 2– 68 kΩ, with the largest errors above 10 kΩ, which would bias the in vivo values low. Second, the 450 µs measurement pulse is longer than typical DBS pulses, so the polarization it includes would bias the values high relative to the load seen by shorter pulses. Because these biases act in opposite directions, the direction of the net error is unknown, and the values below are best treated as an estimate of the load range rather than a calibrated impedance.

### 3.3 In vivo impedance and the compliance envelope

Across the five electrodes, impedance ranged from 32 to 80 kΩ (49.1 ± 9.9 kΩ, mean ± SD over all electrode-sessions). The weekly mean was 43.0 ± 7.4 kΩ at 1 week, peaked at 54.2 ± 14.8 kΩ at 3 weeks and returned to 44.6 ± 7.5 kΩ by 8 weeks, remaining within 43–54 kΩ throughout (Figure 4A). Mean impedance of individual electrodes ranged from 42.7 to 64.2 kΩ, so differences between electrodes (about 1.5-fold) were larger than the change in the group mean over time.

**Figure 4.**
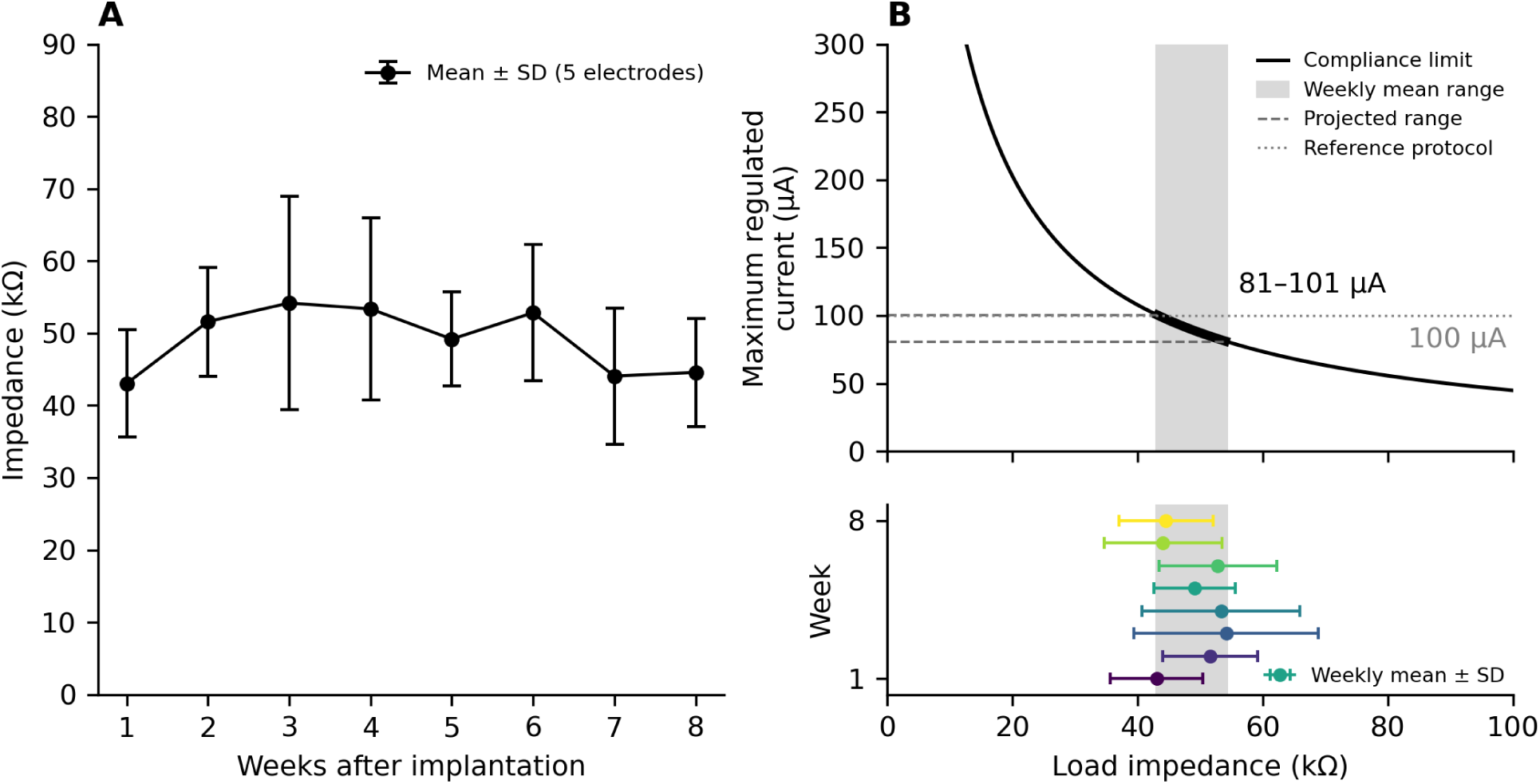
In vivo electrode impedance and the compliance envelope. *(A)* Impedance of implanted platinum electrodes from 1 to 8 weeks after implantation, a pulse-derived estimate of the effective load computed as pulse voltage over delivered current with 10 µA, 450 µs pulses (five readings averaged per electrode and session). Points show the mean ± SD across five electrodes from three mice; three of eight implanted electrodes were excluded (Section 3.2). *(B)* Top: largest regulated current of FLEX-DBS as a function of load impedance, *I*_max_ = 4.6 V/(*|Z|* + 2.66 kΩ), with the swing measured into 47 kΩ. The gray band spans the range of weekly mean impedances in (A), 43–54 kΩ, and the dashed lines project it to the corresponding current range, 81–101 µA; the dotted line marks the 100 µA reference protocol. Bottom: weekly mean ± SD impedance on the same impedance axis, colored from week 1 (bottom) to week 8 (top).

These loads place FLEX-DBS in its compliance-limited regime. With an output swing of about 4.6 V, measured into 47 kΩ, and 2.66 kΩ of sense and filter resistance in series with the electrode, the largest regulated current is approximately *I*_max_ = 4.6 V/(*|Z|* + 2.66 kΩ) (Figure 4B). The weekly mean impedances correspond to 81–101 µA, and individual electrode-sessions to 56–131 µA, compared with the 600 µA nominal range of the application. The 100 µA reference protocol was within compliance in 11 of 40 electrode-sessions, and in 0 to 4 of the 8 sessions for individual electrodes. A battery-direct design with *∼*3 V of compliance would reach only *∼*38–93 µA into the same loads. The usable amplitude is set by electrode impedance and compliance together, not by the programmable range, and it differs between electrodes more than it changes over time after implantation.

## 4. Electrical validation of the wireless stimulator

### 4.1 Current accuracy and pulse fidelity

Both outputs of a device running firmware v0.5 were loaded with 1 kΩ resistors, and one output was recorded with a Joulescope JS220 Precision Energy Analyzer at 500 kHz for *∼*11 s at amplitude settings of 10, 50 and 100 % (180 µs, 130 Hz; Figure 5). The same three settings were then recorded with both outputs loaded with nominal 47 kΩ resistors, within the range of effective loads estimated in vivo. Delivered current was taken as the voltage across the load divided by the load resistance. The meter’s own current channel showed 4–18 µs spikes after each edge from automatic range switching, so both channels were passed through a 38 µs median filter, which removes impulses while preserving steps; charge computed from the voltage changed by less than 0.5 nanocoulombs (nC) with filtering.

**Figure 5.**
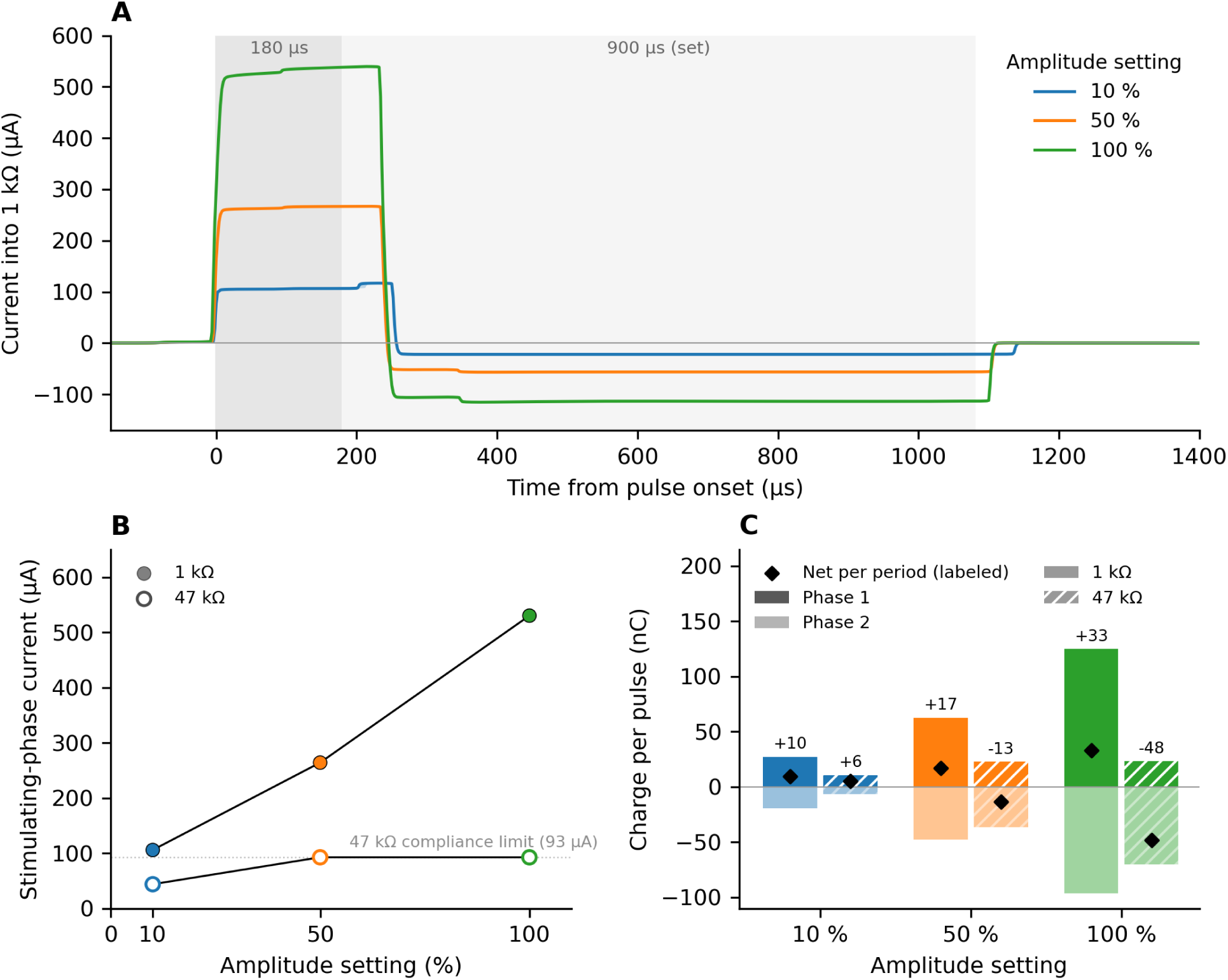
Bench output into 1 kΩ and 47 kΩ loads. Both outputs of a FLEX-DBS device (firmware v0.5) were loaded with 1 kΩ, or with a nominal 47 kΩ within the in vivo load range, and one output was recorded at 500 kHz for *∼*11 s at each of three amplitude settings (10, 50 and 100 %; 180 µs, 130 Hz). Current is the voltage across the load divided by the load resistance, after a 38 µs median filter that removes range-switching artifacts of the meter. *(A)* Median pulse at each setting into 1 kΩ (1545–1602 pulses); the 5th–95th percentile band lies within the line width. Shading marks the set stimulating (180 µs) and recovery (900 µs) phases. *(B)* Mean plateau current of the stimulating phase against the amplitude setting into 1 kΩ (filled) and 47 kΩ (open). The dotted line marks the 92.6 µA compliance limit into 47 kΩ; the 10 % point at 47 kΩ is reduced by slow edges. *(C)* Charge in each phase of the median pulse at 1 kΩ (solid) and 47 kΩ (hatched), and the mean net charge over a full period (labeled; nC, nanocoulombs). At every setting and load the delivered frequency was 142.5 Hz; into 1 kΩ the stimulating phase lasted 238–254 µs and the recovery phase 854–878 µs.

Delivered amplitude increased with the setting and was highly repeatable from pulse to pulse. The stimulating phase measured 106, 264 and 530 µA at 10, 50 and 100 %, with a pulse-to-pulse SD below 0.5 µA at every setting (Figure 5B). A line through the three settings has a slope of 4.7 µA per percent and an intercept of +47 µA, against the 6 µA per percent shown in the application. The application value is therefore a reproducible setting rather than a calibrated current: the device delivers more than the nominal value at low settings and less at high settings. The recovery phase scaled with the stimulating phase at 1:4.7–4.9 (22, 56 and 113 µA).

Pulse timing differed from the set values by a similar amount at every amplitude. At the 180 µs setting the stimulating phase lasted 238–254 µs and the recovery phase 854–878 µs, against 900 µs set, so the two phases did not carry equal charge: the median pulse delivered 27, 63 and 125 nC in the stimulating phase and 19, 48 and 96 nC in the recovery phase, leaving a net 9.5, 17.3 and 33.0 nC per period in the stimulating direction (26–35 % of the stimulating-phase charge; Figure 5C). The 130 Hz setting delivered 142.5 Hz at every amplitude, a 7.02 ms period, because firmware v0.5 sets the period in whole milliseconds (Table 2), and the period varied with an SD of 29–38 µs, about one 30.5 µs timer tick.

Into 47 kΩ, delivered current was limited by compliance (Figure 5B). At the 50 and 100 % settings the stimulating phase was flat-topped at 92.6 µA, which implies an output swing of 4.6 V across the load and the 2.66 kΩ of sense and filter resistance in series with it. The recovery phase reached 47 µA at 50 %, below the 56 µA delivered into 1 kΩ, and was also limited at 100 %, at about 88 µA (4.4 V). At 10 %, below the limit, the edges were slow at this load: the stimulating phase was still rising when it ended (peak 51 µA, against 106 µA into 1 kΩ), and the cause, possibly capacitance on the load side, was not identified. Because these effects reduce the stimulating phase more than the recovery phase, the net charge per period changed sign with amplitude, at +5.5, *−*13.3 and *−*48.4 nC for 10, 50 and 100 % (Figure 5C). At a 0 % setting into 47 kΩ the output carried a residual of about 1 nC per period during the active window, small relative to the imbalance above; in use, stimulation is turned off rather than set to 0 %. Other pulse widths and loads were not characterized.

### 4.2 Battery and operating duration

Battery current was measured at the cell terminals in each operating state with the same Joulescope JS220 (Table 3). The device draws 1.47 mA when idle and not advertising, 1.62 mA while advertising, 1.89 mA while connected to the application, and 3.15–3.42 mA while stimulating at amplitude settings labeled 60 and 400 µA, respectively, into 1 kΩ with a 180 µs pulse width at the 130 Hz frequency setting (142.5 Hz delivered).

**Table 3.** Measured battery current and projected runtime on two size-13 zinc-air cells in series (*∼*280 mAh; 90 % of nameplate capacity assumed usable). Stimulation rows use a 1 kΩ load, a 180 µs pulse width and the 130 Hz setting, which delivers 142.5 Hz, and give the amplitude as labeled in the application. The continuous-stimulation projection was confirmed in a manual bench test that was not logged.

| Operating state | Battery current | Projected runtime |
| --- | --- | --- |
| Idle, not advertising | 1.47 mA | ∼7.1 d |
| Advertising | 1.62 mA | ∼6.5 d |
| Connected to app | 1.89 mA | ∼5.6 d |
| Stimulating, 60 μA | 3.15 mA | ∼3.3 d |
| Stimulating, 400 μA | 3.42 mA | ∼3.1 d |

Two features of these numbers define the usable experimental window. First, raising the amplitude setting from 60 to 400 µA increased battery current by only 0.27 mA, so delivered amplitude contributes relatively little to the power budget. Entering the stimulation state adds 1.68–1.95 mA over idle; this reflects circuitry and processing that are active while the stimulation protocol runs, and how it divides among subsystems was not measured. Second, even at the highest measured current a fresh pair of cells has a projected runtime of about three days of continuous stimulation, and more than five days of standby with the phone connected. The power budget therefore permits hours-long, overnight and multiday sessions without disturbing the animal, with longer studies possible through a brief battery exchange, which requires detaching the device and pauses stimulation; charge balance in the present firmware is a separate constraint. Zinc-air cells are inexpensive and widely available, which suits a device whose cells are replaced frequently.

The device is intended for chronic studies with scheduled battery exchange, not as a fully implanted, maintenance-free system.

## 5. Comparison with existing methods

Table 4 places FLEX-DBS alongside published wireless rodent stimulators that are small enough for mice or are architecturally instructive. The columns are the properties that most often determine whether a system can run a given experiment: species and mass, channels, output type, supply or compliance, waveform and stimulation modes, how parameters are set, and power and operating duration. Supplementary Table S7 adds programmable ranges, dimensions and release status.

**Table 4.** FLEX-DBS in the context of published wireless rodent stimulators. Values as reported by the authors; blank, not reported; parentheses, an alternate value from the same source. Ranges, dimensions and release status: Table S7. CC, constant current; CV, constant voltage.

| System | Species tested | Mass with battery | Channels | Output | Supply / compliance | Waveform and modes | Parameter control | Current draw; operating duration |
| --- | --- | --- | --- | --- | --- | --- | --- | --- |
| **FLEX-DBS (this work)** | Mouse | 3.71 g with shroud (two zinc-air cells) | **2; shared amplitude and timing<sup>†</sup>** | CC | ±5 V rails; ~4.6 V swing (measured) | Asymmetric biphasic, 1:5 recovery; **continuous or burst<sup>†</sup>** | **Real-time BLE from an iOS app; autonomous after disconnect<sup>†</sup>** | 3.15–3.42 mA stimulating; ~3 d per cell pair (projected) |
| t-IPG, Paulat et al. (2026) | Mouse (s-IPG also in rats and mice) | 0.9 g (s-IPG 1.6 g) | 2, time-delayed | CC | 3 V lithium cell, direct | Cathodic phase with long low-amplitude anodic phase; theta- and beta-burst modes | Predefined paradigms via near-field transponder | 21.3 µA maximum; 49 d voltage-compliant |
| STELLA, Plocksties et al. (2021) | Rat (proposed for mice >20 g) | 4.0 g encapsulated (CR1225); 2.6 g (CR1216) | 2, synchronized | CC | Boost 3.1–3.7 V; 2.2–2.8 V at electrode | Monophasic, passive charge balance | Firmware set before implantation; Hall sensor steps through presets | 7.6–8.3 µA; ≥12 wk in vitro, 42 d in vivo |
| Grotemeyer et al. (2024) | Mouse | 2 g | 1 | CC | 3 V (two 1.5 V cells) | Monophasic, passive discharge | Cable programmer, reprogrammable after implantation; reed switch on/off | ~20 µA; 52 d in vitro |
| Fleischer et al. (2020) | Mouse | 2.8 g | 1 | CC | 3 V (two V364 cells) | Monophasic, passive discharge | Programmed before mounting; switch on/off | ~20 µA; ~30 d in vitro |
| de Haas et al. (2012) | Mouse | 2.1 g with batteries and lead | 2, alternating pulses | CC (adaptive resistance) | 4.65 V (three 337 silver-oxide cells); regulates into 15–45 kΩ | Symmetric biphasic | Amplitude set by reprogramming in a fixture; magnet on/off | 15 d standby, 10 h stimulating |
| Alpaugh et al. (2019) | Mouse | 4.7 g | 1 | CC | 3 V (two 393 cells); 2.1 V output headroom | Balanced biphasic | Preprogrammed via external circuit; no radio | ~2 wk per cell set, ≥30 d with weekly exchange |
| Kouzani et al. (2013) | Rat (designed for mice) | 3.27 g (head-mount) | 1 | CC | 3.1 V (two silver-oxide cells) | Monophasic, capacitor-coupled | Preprogrammed; changes require reflashing or a resistor change | ~870 µA; 12.3 d on bench, 7 d in vivo |
| Pinnell et al. (2018) | Rat | 2.8 g with battery and head cap (text: 2.7 g) | 2 | CC | 12.29 V regulated | Monophasic or biphasic; dual-frequency burst | Wired programmer; amplitude by potentiometer; magnet on/off | 35 µA in low-power state; ~30 h stimulating |
| Burton et al. (2021) | Rat | 0.046 g, battery-free | 1 bipolar electrode | CV | 1.5–5.5 V (regulated) | Biphasic | Real time via modulation of RF power | 9.35 mW while stimulating; >36 d in vivo with RF power |
| Shen et al. (2026) | Rat | 12.5 g (7.5 g battery) | Not stated; used unilaterally | CV | 3.7 V Li-ion | Biphasic | Real-time BLE (MATLAB interface) | ≥300 h on 400 mAh |

<sup>†</sup>New paradigm enabling.

Three patterns emerge. First, the mouse-scale systems with runtimes of weeks (Paulat et al., 2026; Plocksties et al., 2021; Grotemeyer et al., 2024; Fleischer et al., 2020) run from supplies of about 3 V, either directly from the cells or through a small boost that leaves 2.2–2.8 V at the electrode, and draw roughly 8–22 µA. Their parameters are set before implantation or through a near-field or cable programmer, and a magnet or reed switch turns stimulation on and off or steps through presets. Their average currents are two orders of magnitude below the 3.15–3.42 mA of FLEX-DBS, a gap that delivered amplitude does not explain and that reflects the circuitry FLEX-DBS keeps active to provide compliance rails, precision analog stages and a radio. The low supply also bounds the load these systems can drive: Alpaugh et al. (2019) note that their 2.1 V of output headroom delivers the full 149 µA only into loads below about 20 kΩ, well below the 32–80 kΩ measured in vivo here. Second, the systems that offer real-time wireless control are rat-scale or externally powered and regulate voltage rather than current: Burton et al. (2021) harvest power from a radio-frequency field in the arena, and Shen et al. (2026) use BLE with a 7.5 g battery. Pinnell et al. (2018) combine constant current, two channels and 12.29 V compliance in a 2.8 g package, but set parameters through a wired programmer and potentiometers and run for about 30 h. Third, no system compared here combines bilateral constant-current output, compliance rails of ±5 V, continuously programmable amplitude, pulse width and frequency, burst mode, and real-time phone control. That combination is what FLEX-DBS contributes, and it is bought with runtime measured in days rather than weeks.

For a fully implanted study with fixed parameters and no routine access to the animal, an architecture such as that of Paulat et al. (2026) or Plocksties et al. (2021) is more suitable; FLEX-DBS targets experiments in which parameters must be changed at the cage.

## 6. In vivo functional validation

### 6.1 Experimental paradigm

In our rodent DBS protocols, stimulation amplitude is titrated in each animal against an observable effect, and the amplitude at which side effects such as muscle twitches appear is a natural upper reference. A preclinical DBS system should therefore be able to deliver amplitudes in the therapeutic range and the larger amplitudes at which side effects appear. DBS of the cingulate cortex is under active investigation for chronic pain that is refractory to conventional medical management (Boccard et al., 2017; Shirvalkar et al., 2025). We hypothesized that high-amplitude cingulate stimulation would produce freezing or prolonged immobility. We therefore delivered high amplitude settings intended to evoke a robust stimulation-associated behavioral phenotype in vivo.

Six mice implanted with cingulate DBS electrodes (Section 3.1), which were being studied in a separate chronic-pain paradigm, had also received a unilateral chronic constriction injury (CCI) of the sciatic nerve, adapted from Bennett and Xie (1988). Briefly, the CCI was performed by making a small incision on the thigh, exposing the sciatic nerve by blunt dissection between the gluteus maximus and biceps femoris, and tightly ligating the sciatic trunk with two chromic gut sutures. Mice were placed in an open-field chamber (30 cm × 30 cm inner dimensions) and allowed to acclimate for 5 minutes before behavioral recording. Position was tracked with Bonsai (Lopes et al., 2015) using a webcam mounted above the chamber (Figure 6A). Each mouse was recorded once, in a single session of 17.5–20.7 minutes comprising a baseline period, a period of bilateral DBS at high amplitude settings and a post-stimulation period. DBS was switched on 5.5–6.9 minutes and off 11.7–14.4 minutes after the start of recording. These mice were separate from those used for the impedance measurements in Section 3.2. Stimulation was delivered by FLEX-DBS with firmware v0.5, as in the rest of this paper, at an amplitude setting above 100 µA, 450 µs pulse duration and 130 Hz pulse frequency (the 130 Hz setting is delivered as 142.5 Hz, Table 2). Electrode impedance and delivered current were not measured in these mice; at the electrode loads measured in Section 3.2, the delivered current would be limited by compliance to approximately 81–101 µA (Figure 4B).

**Figure 6.**
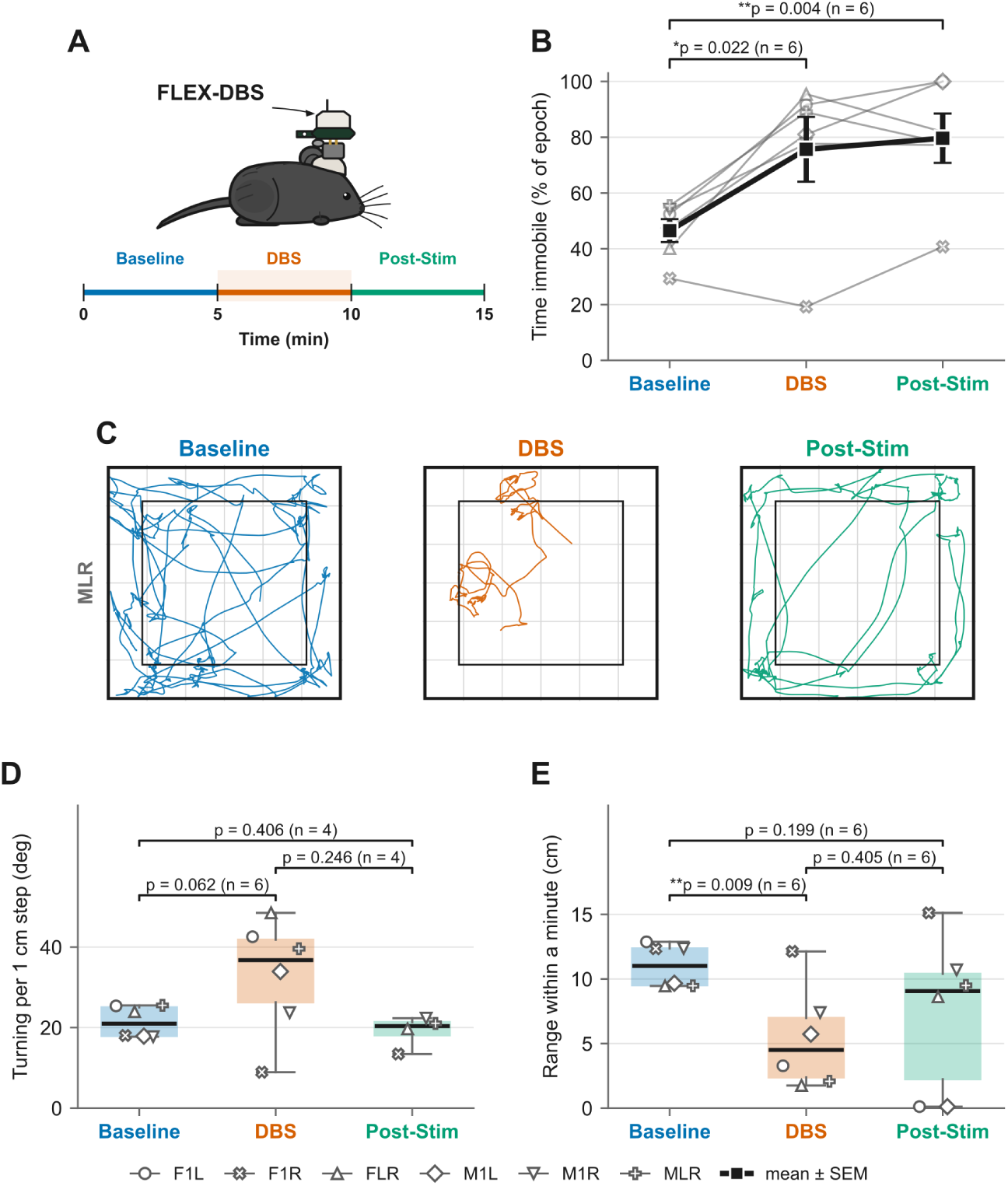
Open-field behavior before, during and after FLEX-DBS at high amplitude settings. *(A)* Recording setup and approximate session timeline (actual timing in Section 6.1). Period colors are used throughout. *(B)* Percent of each analyzed period spent immobile. Gray lines and open markers show individual animals (n = 6), with marker shape identifying each animal in B, D and E; black squares show mean ± SEM. *(C)* Trajectories of one example animal (MLR) in each period. The outer square marks the extent of tracked positions, and the inner square the equal-area center zone. *(D)* Mean absolute change in heading between consecutive 1 cm steps of path; undefined for animals that did not move. *(E)* Locality, the radius of gyration of position within 1 min bins, averaged over the period. In D and E, boxes show median and interquartile range, and whiskers extend to the most extreme values within 1.5 × the interquartile range. Brackets give two-tailed paired t-tests with the number of animals in each comparison; *p < 0.05, **p < 0.01.

### 6.2 Video-based behavioral analysis

The position of each mouse in the open-field chamber was tracked throughout the baseline, DBS and post-stimulation periods (Figure 6C). Video was saved at 320 × 240 px (MPEG-4), and the centroid of the animal was tracked online in an 894 × 877 px tracking frame, with each position exported with its acquisition timestamp. Frame timing was irregular (median approximately 31 frames/s; inter-frame intervals 13–58 ms), so speed was computed as displacement divided by the measured interval between frames, and all averages were weighted by time. Pixel-to-centimeter calibration was measured separately for each recording from its own video, as the wall-to-wall span of the arena floor detected in a median background image (27.30–27.73 px/cm across recordings), because the camera position shifted slightly between sessions. Trajectories were preprocessed in the following order: (1) tracking gaps of up to 0.25 s were linearly interpolated, and longer gaps were left missing; (2) frames implying a speed above 100 cm/s, which corresponded to the tracker jumping to the experimenter’s hand, were rejected iteratively and replaced by linear interpolation; (3) a 5-frame median filter was applied; and (4) position was smoothed with a Gaussian filter (standard deviation 0.12 s, applied on the frame index at the median frame rate).

The baseline, DBS and post-stimulation periods were separated by brief handling events in which the experimenter’s hand entered the chamber to switch stimulation on and off. Handling events were detected automatically in the video, and detections less than 60 s apart were merged into a single event. Because handling transiently increased locomotion, every analysis window began 90 s after the preceding handling event. This applied to all three periods: the baseline window began 90 s after the start of recording, the DBS window 90 s after the stimulation-on event, and the post-stimulation window 90 s after the stimulation-off event. The 90 s buffer was fixed before the second group of animals was recorded and was not adjusted afterward. Baseline windows ended at the stimulation-on event, DBS windows at the stimulation-off event, and post-stimulation windows at the end of recording. Analyzed durations were 241–324 s (baseline), 263–339 s (DBS) and 225–288 s (post-stimulation).

Immobility was defined by classifying each inter-frame interval as mobile or immobile using speed thresholds with hysteresis. A mobile bout began when speed exceeded 2 cm/s and ended when speed fell below 1 cm/s, with minimum bout durations of 0.5 s (mobile) and 1.0 s (immobile). Percent time immobile was the time-weighted fraction of the analysis window not classified as mobile. The thresholds, the 100 cm/s speed ceiling and the smoothing parameters were chosen values rather than values taken from the literature. Turning was quantified after rediscretizing each trajectory into steps of constant 1 cm length, independent of speed or elapsed time (Bovet and Benhamou, 1988). A new point was recorded each time the animal had moved 1 cm in a straight line from the previous point, and paths were split at untracked gaps so that no step spanned missing data. The turning angle was the change in heading between consecutive steps, wrapped to ±180 degrees. Turning for each period was the mean absolute turning angle, which counts turns in either direction, and required at least 20 steps (20 cm of path); otherwise it was left undefined. Locality was quantified as the radius of gyration, the time-weighted root-mean-square distance of the animal’s position from its mean position. It was computed in consecutive 60 s bins from the start of each analysis window, with bins containing less than 30 s of valid tracking discarded, and averaged across bins weighted by tracked time.

Periods were compared with two-tailed paired t-tests using the animal as the unit of analysis (n = 6). Animals without a defined value in a period were excluded only from comparisons involving that period. For immobility, a Wilcoxon signed-rank test was also computed for baseline versus DBS. To estimate the contribution of within-session habituation, immobility during DBS was additionally compared with the final 2 min of the baseline window. Turning and locality were exploratory analyses developed after the immobility result, and their p-values are reported without correction for multiple comparisons. Analyses were performed in Python; turning, locality, statistics and figures were computed with Python 3.13.5, NumPy 2.5.3, pandas 3.0.6, SciPy 1.18.1 and Matplotlib 3.11.2.

### 6.3 Immobility increased during stimulation at high amplitude settings

To confirm that the device delivered stimulation sufficient to produce observable motor effects, mice were recorded in an open field before, during and after FLEX-DBS at high amplitude settings in a single session (Figure 6A). Across the six animals, the percentage of time spent immobile was 46.5 ± 4.1 % during baseline, 75.7 ± 11.6 % during DBS and 79.7 ± 8.9 % after stimulation (mean ± SEM; Figure 6B), a 62.7 % increase in the group mean from baseline to DBS. Immobility was higher during DBS than at baseline (paired t-test, t(5) = −3.26, p = 0.022) and remained higher after stimulation ended (t(5) = −4.95, p = 0.004). Five of six animals showed the increase; the exception, F1R, became less immobile during DBS (29.4 % to 19.2 %), and a Wilcoxon signed-rank test of baseline versus DBS gave p = 0.0625. Because all periods were recorded in one session without a sham control, this increase is associated with stimulation but cannot be separated from within-session habituation. When DBS was compared only with the final 2 min of baseline (50.4 ± 7.3 % immobile), the difference was smaller and not significant (t(5) = −2.34, p = 0.066), and immobility did not return toward baseline levels after stimulation ended.

In exploratory analyses that were not corrected for multiple comparisons, movement during DBS was more tortuous and more spatially confined than at baseline (Figure 6C–E). Turning per 1 cm step rose from 21.4 ± 1.6 to 32.9 ± 5.9 degrees (t(5) = −2.40, p = 0.062), increasing in five of six animals, again with the exception of F1R. The within-minute radius of gyration fell from 11.0 ± 0.7 to 5.4 ± 1.6 cm (t(5) = 4.13, p = 0.009). Neither measure differed significantly between DBS and post-stimulation or between baseline and post-stimulation: for turning, n = 4 because two animals did not move after stimulation, with p = 0.246 and p = 0.406; for locality, n = 6, with p = 0.405 and p = 0.199. Locality partly reflects immobility, because a stationary animal scores near zero. Together, these data show motor changes during FLEX-DBS at high amplitude settings, consistent with effective stimulation through implanted electrodes. They do not establish that stimulation caused the changes or that the changes reversed when stimulation ended.

## 7. Discussion

FLEX-DBS provides flexible, bilateral, constant-current DBS with real-time wireless control for experiments in freely moving rodents. Its measured power consumption permits a projected three days of continuous stimulation per pair of cells, with longer studies possible through a brief battery exchange. Removing the tether while retaining the ability to set, change and verify parameters at the cage allows paradigms our laboratory has used with tethered hardware (Creed et al., 2011, 2012, 2013; Aleksandrova et al., 2013) to move into home-cage, social and extended-duration assays in which a tether is a significant confound.

The in vivo load measurements reinforce that DBS delivery is a systems problem. Electrode impedance and stimulator compliance jointly determine whether the commanded constant current is actually delivered, and impedance differs between electrodes and varies over the weeks after implantation. A programmable current range alone therefore does not fully characterize a stimulator. For the implanted platinum electrodes used here, with pulse-derived effective loads of 32–80 kΩ and weekly means of 43–54 kΩ, the ±5 V rails of FLEX-DBS support roughly 56–131 µA, well below the 600 µA programmable range and centered on the 100 µA protocols used in prior work. We therefore recommend reporting the compliance limit alongside the measured load distribution (Figure 4B) for any rodent stimulator.

Two practical consequences follow. First, electrode impedance is the most direct lever on usable amplitude, and future protocols will establish a method for fabricating lower-impedance electrodes. Second, amplitude should be titrated in each animal against its behavioral effect, as in our protocols, rather than taken from the application’s nominal value. A configurable stimulator lets the experimenter select an amplitude that is deliverable into the implanted load and behaviorally effective, and adjust it if either condition fails.

Flexible multiday systems and fully implanted long-duration systems occupy different optimization spaces, and the measured current budget shows why. Raising the amplitude setting from 60 to 400 µA adds only 0.27 mA, whereas entering the stimulation state adds 1.68–1.95 mA over idle, so most of the current drawn during stimulation reflects circuitry and processing required while the protocol runs rather than delivered amplitude. The design carries that overhead deliberately, because each block enables a capability of FLEX-DBS: a boost converter and inverter provide ±5 V compliance; a continuously powered precision DAC and reference set the current reproducibly; an awake microcontroller sequences each pulse; and the radio stack permits configuration and verification at the cage. Power was reduced within these constraints, most visibly by shutting the output amplifier down between pulses, but maintenance-free operation for weeks or months was not the primary objective.

A fully implanted derivative would invert these priorities: battery-direct operation at *∼*3 V if electrode impedance allows it, a shared current source with hardware-timed polarity switching, asymmetric or passive charge recovery, a radio used only to configure, and a microcontroller whose hardware peripherals can sequence a pulse without waking the core. Published mouse-scale systems show that this combination reaches average currents of roughly 8–22 µA (Paulat et al., 2026; Plocksties et al., 2021; Grotemeyer et al., 2024; Fleischer et al., 2020).

## Limitations

Firmware v0.5 delivers the 130 Hz setting as 142.5 Hz and leaves a net charge of 26–35 % of the stimulating-phase charge per pulse into 1 kΩ; this must be corrected before prolonged in vivo stimulation, and it applies to the stimulation delivered in Section 6. Below compliance, the finite output impedance of the Howland pumps (about 0.9 MΩ worst case with 0.1 % resistors; Section 2.3) lets delivered current fall by up to 3–8 % into the loads measured here, so amplitude settings should be read against the measured load and the bench measurements rather than as absolute currents. The two channels share amplitude and timing, and the 80 Hz frequency floor excludes the low-frequency protocols of comparative and therapeutic interest (Creed et al., 2011, 2015). Finally, the bench measurements cover one pulse width into two loads, the impedance data come from five electrodes in three mice, and the in vivo test was a single session in six mice without a sham control, so the immobility increase cannot be separated from within-session habituation.

## Future development

The firmware, application and protocol are released, and the schematic is available on request, so that others can reproduce or adapt the platform (Supplementary material). Several extensions are firmware-level within the present hardware: computing the stimulation period in timer ticks rather than whole milliseconds, which would deliver 130 Hz and other frequencies exactly; calibrating the phase-timing compensation and recovery amplitude against bench measurements so that the two phases carry equal charge; independent per-channel timing; a wider frequency range; a selectable recovery ratio; and over-the-air firmware update. At the hardware level, on-board sensing of delivered current or electrode voltage would report the stimulation actually delivered and track electrode impedance over the course of an experiment, supporting amplitude titration in each animal.

## 8. Conclusions

FLEX-DBS is a head-mounted, bilateral, constant-current stimulator with real-time smartphone control, intended for experiments in which parameters must be set, changed and verified at the cage. Its measured power consumption permits a projected three days of continuous operation per cell pair; firmware v0.5 requires improved charge balancing before prolonged in vivo stimulation. Its ±5 V compliance rails deliver roughly 56–131 µA into implanted platinum electrodes with pulse-derived effective loads of 32–80 kΩ, so the electrode load, not the programmable range, sets usable amplitude in vivo. In vivo, stimulation at high amplitude settings was associated with increased immobility in an open field, consistent with effective stimulation through implanted electrodes.

## Supplementary material

- **S1.** Circuit schematic of the FLEX-DBS board (PDF); available from the corresponding author on request.
- **S2.** Firmware for the BGM220S (Silicon Labs Simplicity Studio project), v0.5. Repository: https://github.com/Neurotech-Hub/Creed-DBS-SiLabs
- **S3.** MouseCap iOS application source. Repository: https://github.com/Neurotech-Hub/MouseCap-iOS
- **S4.** BLE protocol specification (`MCU_BLE_PROTOCOL.md` in S3): service and characteristic UUIDs, command grammar, parameter keys and ranges, and response format.
- **S5.** Shroud design (Fusion 360): https://a360.co/3TqHAxf
- **S6.** Electrode holder design (Fusion 360): https://a360.co/4rEBmq2
- **S7.** Extended comparison table (Table S7, below): programmable ranges, dimensions and release status for the systems in Table 4.

**Table S7.** Extended comparison of FLEX-DBS with published wireless rodent stimulators (full version of Table 4). Values for published systems are as reported by the authors; blank cells, and sizes omitted from the mass column, are not reported in the source. Where a source gives two values, the second is shown in parentheses with its location. CC, constant current; CV, constant voltage.

| System | Species tested | Mass with battery; size | Channels | Output | Supply / compliance | Amplitude, pulse width, frequency; waveform | Parameter control | Current draw; operating duration | Open release |
| --- | --- | --- | --- | --- | --- | --- | --- | --- | --- |
| **FLEX-DBS (this work)** | Mouse | 3.71 g with shroud (two zinc-air cells); 19.5 × 22.5 × 8.72 mm | **2; shared amplitude and timing<sup>†</sup>** | CC | ±5 V rails; ~4.6 V swing (measured) | 0–625 µA ideal (compliance-limited in vivo); 90–600 µs set; ~83–167 Hz delivered; asymmetric biphasic, 1:5 recovery; **continuous or burst<sup>†</sup>** | **Real-time BLE from an iOS app; autonomous after disconnect<sup>†</sup>** | 3.15–3.42 mA stimulating; ~3 d per cell pair (projected) | Firmware, app, protocol (S2–S4); schematic on request (S1) |
| t-IPG, Paulat et al. (2026) | Mouse (s-IPG also in rats and mice) | 0.9 g (s-IPG 1.6 g); 5.3 mm thick | 2, time-delayed | CC | 3 V lithium cell, direct | 1–100 µA, 50–700 µs, 2–257 Hz; cathodic phase with long low-amplitude anodic phase; theta- and beta-burst modes | Predefined paradigms via near-field transponder | 21.3 µA maximum; 49 d voltage-compliant | Partial (GitHub) |
| STELLA, Plocksties et al. (2021) | Rat (proposed for mice >20 g) | 4.0 g encapsulated (CR1225), Ø17.0 × 9.0 mm; 2.6 g (CR1216), Ø15.5 × 7.5 mm | 2, synchronized | CC | Boost 3.1–3.7 V; 2.2–2.8 V at electrode | 42–100 µA, 1 µs–100 % duty, 10–5000 Hz; monophasic, passive charge balance | Firmware set before implantation; Hall sensor steps through presets | 7.6–8.3 µA; ≥12 wk in vitro, 42 d in vivo | Yes, open license (GitHub) |
| Grotemeyer et al. (2024) | Mouse | 2 g; 18 × 16 × 5 mm with housing | 1 | CC | 3 V (two 1.5 V cells) | 10–500 µA, 60–500 µs, 10–200 Hz; monophasic, passive discharge | Cable programmer, reprogrammable after implantation; reed switch on/off | ~20 µA; 52 d in vitro | Analysis code only; commercialized |
| Fleischer et al. (2020) | Mouse | 2.8 g; board 9 × 9 × 1 mm (abstract: 10 × 10 mm) | 1 | CC | 3 V (two V364 cells) | 0–500 µA (text: up to 300 µA), 60–500 µs, 10–200 Hz (text: 10–500 Hz); monophasic, passive discharge | Programmed before mounting; switch on/off | ~20 µA; ~30 d in vitro |  |
| de Haas et al. (2012) | Mouse | 2.1 g with batteries and lead; Ø8 × 30 mm | 2 (bilateral), pulses alternate between channels | CC (adaptive series resistance) | 4.65 V (three 337 silver-oxide cells); regulates into 15–45 kΩ | 20–100 µA, 60 µs, 131 Hz fixed; symmetric biphasic | Amplitude set by reprogramming in a fixture (code table and resistor changes); magnet on/off; IR LED status | 15 d standby, 10 h stimulating |  |
| Alpaugh et al. (2019) | Mouse | 4.7 g | 1 | CC | 3 V (two 393 cells); 2.1 V output headroom | Fixed 149 µA, ~103 µs, ~130 Hz; balanced biphasic | Preprogrammed via external circuit; no radio | ~2 wk per cell set, ≥30 d with weekly exchange | Component list only |
| Kouzani et al. (2013) | Rat (designed for mice) | 3.27 g (head-mount) | 1 | CC | 3.1 V (two silver-oxide cells) | Fixed 200 µA, 90 µs, 130 Hz; monophasic, capacitor-coupled | Preprogrammed; changes require reflashing or a resistor change | ~870 µA; 12.3 d on bench, 7 d in vivo | Schematic and pseudocode |
| Pinnell et al. (2018) | Rat | 2.8 g with battery and head cap (text: 2.7 g); device Ø12.5 × 5 mm | 2 | CC | 12.29 V regulated | 20 µA–2 mA, 10 µs–100 % duty, 0.1–5000 Hz; monophasic or biphasic; dual-frequency burst | Wired programmer; amplitude by potentiometer; magnet on/off | 35 µA in low-power state; ~30 h stimulating | Design files and assembly instructions (Figshare) |
| Burton et al. (2021) | Rat | 0.046 g, battery-free | 1 bipolar electrode | CV | 1.5–5.5 V (regulated) | 1.5–5.5 V, ≥20 µs pulses, up to 25 kHz; biphasic | Real time via modulation of RF power | 9.35 mW while stimulating; >36 d in vivo with RF power |  |
| Shen et al. (2026) | Rat | 12.5 g (7.5 g battery); 34 × 42 × 15 mm with housing | Not stated; used unilaterally | CV | 3.7 V Li-ion | 2.3–3 V, 40–340 µs, 80–150 Hz; biphasic | Real-time BLE (MATLAB interface) | ≥300 h on 400 mAh |  |

<sup>†</sup>New paradigm enabling.

## Author contributions

RDG implemented the experiments, performed the impedance bench testing and carried out animal handling and testing. MCC conceived and designed the experimental approach. MG designed the hardware, carried out the technical implementation, and performed the electrical bench testing and analysis. All authors contributed to writing the manuscript.

## Ethics statement

All animal procedures were approved by the Institutional Animal Care and Use Committee at Washington University in St. Louis (protocol 24-0211) and conducted in accordance with the Guide for the Care and Use of Laboratory Animals (NIH).

## Data and code availability

The firmware is publicly available at https://github.com/Neurotech-Hub/Creed-DBS-SiL `abs` (S2), and the iOS application source and BLE protocol specification at https://github.com/N `eurotech-Hub/MouseCap-iOS` (S3, S4). The shroud and electrode holder designs are available at https://a360.co/3TqHAxf and https://a360.co/4rEBmq2. The circuit schematic (S1), the impedance data underlying Figure 4, the bench recordings and analysis scripts underlying Figures 4 and 5, the behavioral tracking data and analysis code underlying Figure 6, and other design files are available from the corresponding author on request.

## Acknowledgments

We thank the Washington University Neurotech Hub for technical assistance with 3D design and hardware fabrication.

